# Analysis of polyclonal and monoclonal antibody to the influenza virus nucleoprotein in different oligomeric states

**DOI:** 10.1101/2024.09.12.612748

**Authors:** Mallory L. Myers, Michael T. Conlon, John R. Gallagher, De’Marcus D. Woolfork, Noah D. Khorrami, William B. Park, Regan K. Stradtman-Carvalho, Audray K. Harris

**Author notes:** To whom correspondence should be addressed; E-mail A.K. H.

## Abstract

Influenza virus nucleoprotein (NP) is one of the most conserved influenza proteins. Both NP antigen and anti-NP antibodies are used as reagents in influenza diagnostic kits, with applications in both clinical practice, and influenza zoonotic surveillance programs. Despite this, studies on the biochemical basis of NP diagnostic serology and NP epitopes are not as developed as for hemagglutinin (HA), the fast-evolving antigen which has been the critical component of current influenza vaccines. Here, we characterized the NP serology of mice, ferret, and human sera and the immunogenic effects of NP antigen presented as different structural complexes. Furthermore, we show that a classical anti-NP mouse mAb HB65 could detect NP in some commercial influenza vaccines. MAb HB65 bound a linear epitope with nanomolar affinity. Our analysis suggests that linear NP epitopes paired with their corresponding characterized detection antibodies could aid in designing and improving diagnostic technologies for influenza virus.

## 1. INTRODUCTION

Influenza virus is a pleomorphic, enveloped virus with a viral membrane which displays viral glycoproteins on its surface, including hemagglutinin (HA), neuraminidase (NA) and Matrix (M2) (1-4). On the interior side of the viral membrane is a layer of the matrix protein (M1) (Harris et al., 2006; Harris et al., 2013). Centrally packaged within influenza virions are segmented genomic ribonucleoprotein complexes (RNPs) (Harris et al., 2006). RNPs are double-helical filaments composed of negative-sense genomic viral RNAs that are bound by molecules of the nucleoprotein (NP) (Arranz et al., 2012; Chenavier et al., 2023; Coloma et al., 2020; Gallagher et al., 2017; Moeller et al., 2012). The structure of the NP monomer subunit is dominated by alpha helical secondary structure, and the tertiary structure can be divided into a head domain, comprised of amino acids 150-250 that forms an offset helical bundle, and the body domain, which is folded from primary sequence amino acids both before and after those forming the head. A C-terminal loop mediates NP-NP oligomerization (Ye et al., 2006). RNPs are associated with a trimeric viral polymerase, and as a combined unit, they are responsible for genomic packaging and replication (Coloma et al., 2020; Moeller et al., 2012). NP is more conserved than HA between the different influenza subtypes. Bioinformatics of over 22,000 sequences found that 95% of the length of nucleoprotein was found to be conserved (Babar and Zaidi, 2015). Direct comparison of HA proteins H1 A/California/04/2009 and H3 A/Brisbane/10/2007 yields 42.4% identity and 59.7% similarity between H1 and H3 HA subtypes, the NP sequences from those same strains yield 89.8% identity and 96.6 % similarity.

NP vaccinations result in both T-cell and B-cell (antibody) outcomes with interplay between antibody effector functions and T-cells mediated by NP antigen. For example, NP vaccination can aid in protection via T-cell effector mechanisms (Carragher et al., 2008; Cookenham et al., 2020; Prasad et al., 2001; Vitelli et al., 2013). NP is also important in eliciting both T-cell and antibody responses that contribute to defense against influenza-induced bacterial diseases to aid in the reduction in lung pathology (Haynes et al., 2012). Previous work demonstrated that anti-NP IgG specifically promoted influenza virus clearance in mice by using a mechanism involving both Fc-Receptors and CD8+ cells (LaMere et al., 2011). In addition, mouse NP immune sera could transfer protection against influenza infection to naïve hosts in an antibody-dependent manner (Carragher et al., 2008). Using transgenic mice expressing human anti-NP antibodies, it was observed that non-neutralizing anti-NP antibodies could induce antibody-dependent cellular cytotoxicity or complement-dependent cytotoxicity to aid in abrogation of disease (Fujimoto et al., 2016). Hence, NP antibodies could provide a means of increasing the breadth of protection to different influenza subtypes. However, a further understanding of the biochemistry of NP antibodies is required to begin to understand the roles NP vaccination and anti-NP antibodies might play in immunity to influenza virus.

The relation between NP use as a vaccine antigen to elicit anti-NP antibodies and the use of both NP and anti-NP antibodies in diagnostics bear on each other and should be further studied for several reasons. First, the detection or quantitation of NP is not required for commercial influenza vaccines. To date, the only requirements for vaccine approval are the inclusion of HA protein and demonstration that it can provide protection in animal models and humans (Bouvier and Palese, 2008; Krammer et al., 2015). Different formulations are currently being utilized for commercial vaccine production: inactivated split-subunit (Friedewald, 1944; Gross and Ennis, 1977; Salk et al., 1940; Webster and Laver, 1966), recombinant HA (Cox et al., 2008; McCraw et al., 2016; Treanor et al., 2011), and live-attenuated influenza viruses(Babu et al., 2014; Broadbent et al., 2014; Chen et al., 2014; Jin and Subbarao, 2015; Krammer et al., 2014; Sobhanie et al., 2016), which may lead to differences in inclusion of non-HA proteins, such as NP. Second, concerns raised in previous studies regarding adverse events from vaccines containing NP continue to be a topic of discussion within the field of influenza vaccine research. For example, studies including the adjuvant AS03 that elicited anti-NP-antibodies found the vaccination as a possible cause of autoimmunity and narcolepsy (Ahmed et al., 2014; Ahmed and Steinman, 2016, 2017; Ahmed et al., 2015; Fujimoto et al., 2016; Steinman and Ahmed, 2015). Third, NP and anti-NP antibodies are widely used in diagnostics assays and kits (Ji et al., 2022; Mizuike et al., 2011). The cross-reactivity of NP-reactive antibodies from both humans and animals permits detection of influenza infection from varied influenza strains (LaMere et al., 2011; Vanderven et al., 2016). Last, specific anti-NP antibodies have been elevated through recombinant engineering, including the use of biotinylated single-domain anti-NP antibodies and nanobodies, again primarily for use in diagnostics test and kits for influenza (Du et al., 2019; Ji et al., 2022).

In this study, to more directly define the linkage between NP structure, biomolecular properties, and its role as a bellwether for influenza infection, we characterized the interactions of polyclonal antibodies to influenza nucleoprotein (NP) in sera of infected hosts such as humans, ferrets, and mice. We analyzed the effects of different NP structural arrangements, including NP-rings and NP-filaments, on NP immunogenicity in mice. We show that purified recombinant NP can be used as a diagnostic to detect anti-NP antibodies in sera either in ELISA or immunoblot formats.

Previous work has established mAb HB65 (Yewdell et al., 1981) as an exemplary anti-NP antibody with a wide range of applications (Nicholls et al., 2012; Starick et al., 2006), including a competitive ELISA detection assay for the detection of NP-specific antibodies in sera of ducks, geese, and wild birds (Starick et al., 2006), or immunohistochemical detection of NP in lung tissues from avian, swine and human samples infected with highly pathogenic and pandemic influenza viruses (Nicholls et al., 2012). Passive immunization with mAb HB65 has been reported to suppress inflammatory cytokines and chemokines and provided protection from H1N1-induced secondary pneumococcal disease but did not reduce viral titers or prevent bacterial infection (Haynes et al., 2012). We selected HB65 for further characterization as a representative anti-NP monoclonal antibody (mAb) for these reasons, and because of its commercial availability. We show that polyclonal antibodies and the classical mAb HB65 can detect the presence of NP in viral lysates and that mAb HB65 can detect and quantitate NP in commercial vaccines preparations. This is important because several diagnostics tests for influenza show variation in detection capacity (Balish et al., 2013; Bose et al., 2014). Our work suggests that further engineering both NP and anti-NP antibodies could aid in the development of more effective influenza diagnostics.

## 2. RESULTS

### 2.1. Analysis of post-infection sera of human and ferrets for NP antibodies

To understand the levels of antibodies to NP in sera of humans and ferrets infected with different subtypes of influenza A viruses, sera were analyzed via immunoassays with recombinant NP as the antigen. Sera samples from humans naturally infected with H5N1 (A/Indonesia/5/2005), which were previously classified as having low, medium, and high titers of anti-HA antibodies (IRR), were used to probe for the level of anti-NP antibodies. Antibodies to NP were detected in all human sera groups, and the observed levels of NP antibodies followed the trend of the levels of reported HA antibodies. For example, the high HA titer group also had the highest level of antibodies to NP, as determined by ELISA (Fig. 1A, Fig. S1A). Further analysis of the high titer group sera indicated that anti-NP could detect linear epitopes as judged by immunoblotting (Fig. 1B). The sera reactivity to NP appeared specific in that lysozyme was not reactive as well as non-infected sera used as controls (Fig. 1B, 1C, 1D).

**Fig. 1.**
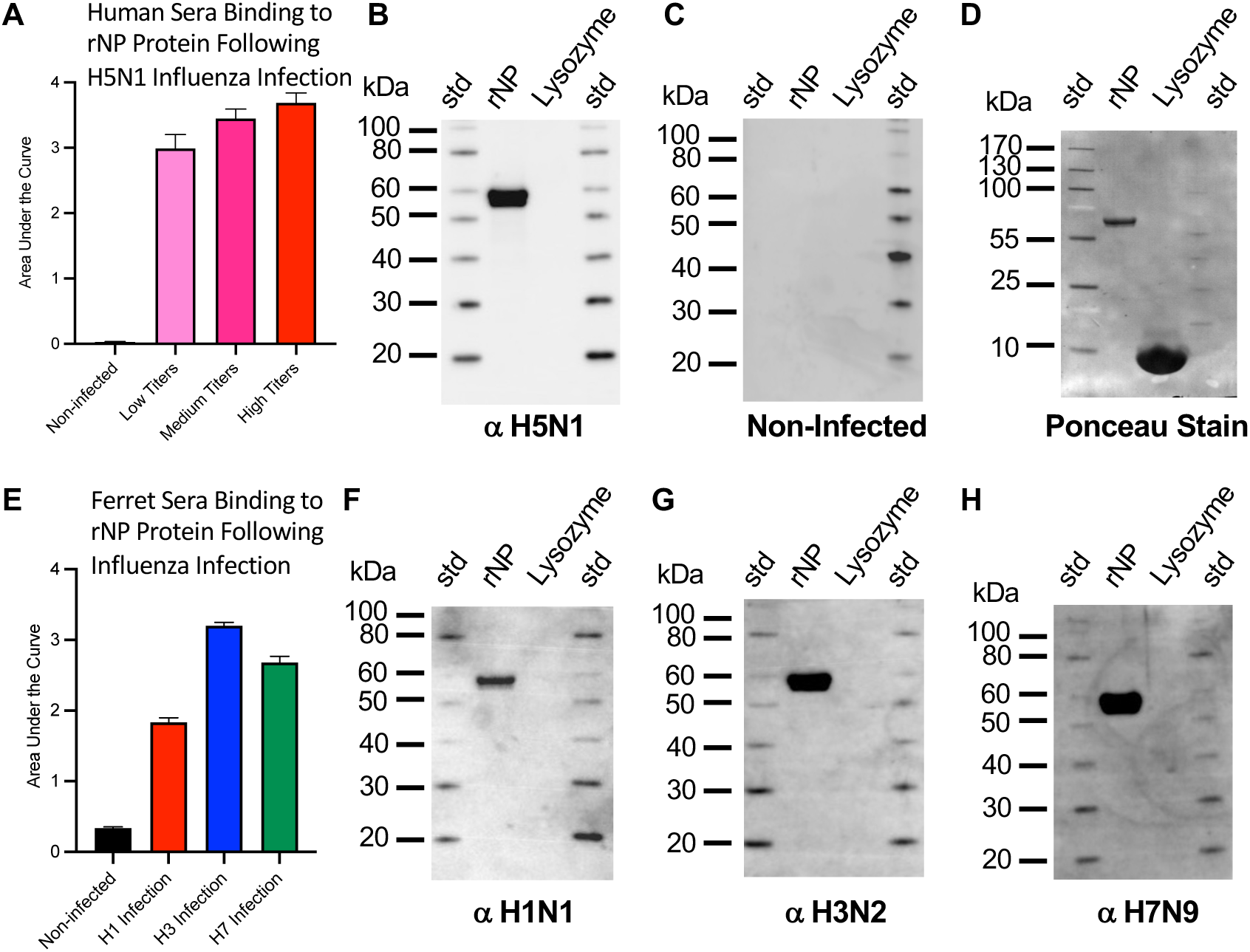
Probing for the presence of anti-nucleoprotein (NP) antibodies in sera of humans and ferrets infected with different subtypes of influenza virus. (A) ELISA analysis of pooled sera from humans infected with H5N1 deemed to have low (pink), medium (magenta), and high (red) titers of virus for detection of antibody binding to rNP. (B) Immunoblot analysis of high titer human sera for detection of rNP. (C) Pooled non-infected sera were used as a negative control and (D) blot from panel C (negative-control) stained with ponceau stain to confirm protein was transferred to the blot. Sera were from human infection with A/Indonesia/5/2005 (H5N1), (E) ELISA analysis of sera of groups of ferrets infected with H1 (red), H3 (blue), or H7 (green) influenza viruses, or non-infected control sera (black), for detection of antibody binding the rNP. (F, G, H) Immunoblot analysis of ferret sera infected with H1, H3, H7 influenza viruses, respectively, for detection of antibody binding to rNP. Sera were from ferret infection with (F) A/California/07/2009 (H1N1), (G) A/Rhode Island/01/2010 (H3N2), and (H) A/Anhui/1/2013 (H7N9), respectively. Immunoblot lanes have molecular weight standards (std) and recombinant NP (rNP) with lysozyme used a negative control. Recombinant NP from influenza A/Brisbane/10/2007 (H3N2) was the antigen used for detection and ferret sera and human sera were used as the primary antibody.

To examine additional strains of influenza virus for their ability to elicit NP antibodies following infection, ferret sera to group 1 (H1N1) and group 2 (H3N2, H7N9) influenza A viruses were probed for binding to rNP via ELISA and immunoblots. The H3N2 infection had higher titers of antibodies to NP than H1N1 and H7N9 infection, as determined by ELISA (Fig. 1E. Fig. S1B). Higher H3N2 reactivity may in part be explained by the rNP used in ELISA being derived from H3N2 virus. Nonetheless, all sera from infected ferrets were able to show binding to rNP in the conformational (Fig. 1E) and linear state (Fig. 1F-H). Sera reactivity to NP appeared specific in that lysozyme was not reactive with infected sera (Fig. 1F, 1G, 1H) and additionally sera from non-infected controls did not react to rNP or lysozyme (Fig. 1C). By immunoblot, NP bands appeared at 55 kilodaltons (kDa) (Fig. 1). Taken together, these results indicated that mammals infected with different subtypes of influenza (e.g. H5N1, H1N1, H3N2, H7N9) elicit cross-reactive antibodies to the conserved NP protein of influenza, and that some of these epitopes are linear epitopes.

### 2.2. Organization and immunogenicity to NP antigen and reactivity of elicited antibodies

The structural organization of recombinant NP protein determines what epitopes may be displayed or occluded for antibody binding, yet it is possible to employ immunoassays such as ELISA without this structural knowledge, leaving a gap in understanding. Thus, to determine the oligomeric state and structural organization of the recombinant NP protein that we used in immunoassays (Fig. 1) we characterized the recombinant NP antigen by electron microscopy. NP was judged to be greater than 95% pure (Fig. 2A) and the NP contained NP oligomers (Fig. 2B). Further image analysis indicated NP oligomers were ring structures of trimers, tetramers, pentamers and hexamers of NP subunits (Fig. 2C panels I-IV, respectively). Separately, NP was isolated from influenza virus (viral NP) in the form of filamentous RNPs (Fig. 2D). To determine if rNP and viral NP were differentially immunogenic, mice were given two intramuscular injections of non-adjuvanted NP preparations at days 0 and 21 (Fig. 2E).

**Fig. 2.**
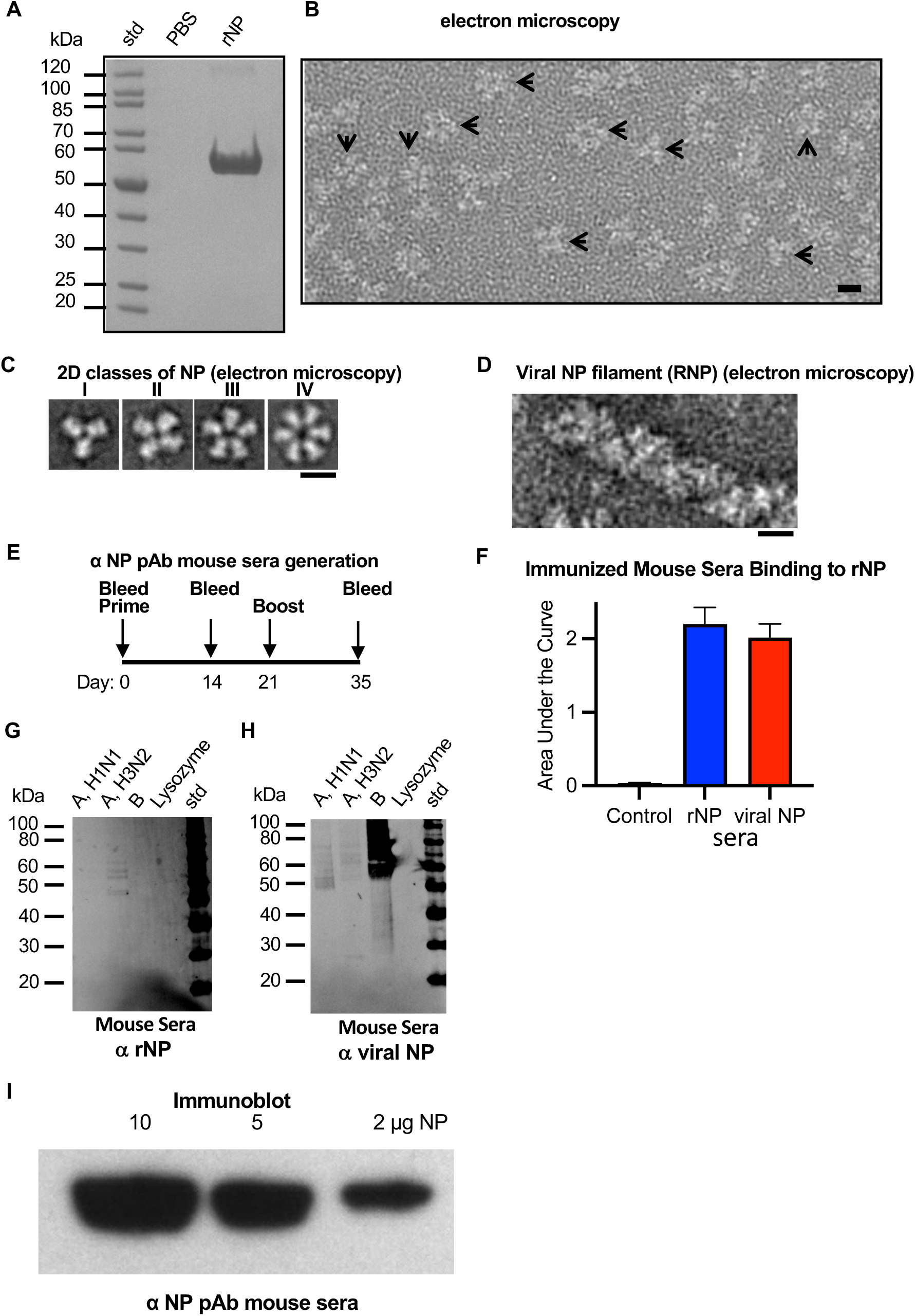
Electron microscopy of recombinant NP and assessment of immunogenicity to recombinant NP and viral NP in mice by immunoassays. (A) Purity of recombinant nucleoprotein (NP) assessed by SDS-PAGE. Lanes were loaded with standards (std) and a PBS spacing control and rNP. (B) Image of a field of NP oligomeric complexes by negative-staining electron microscopy (EM). Arrows denote some complexes. (C) Examples of 2D class averages of NP complexes imaged by EM (panel B). Oligomers are trimers (panel I), tetramers (panel II), pentamers (panel III), and hexamers (panel IV). (D) EM example of viral genomic RNP filament (viral NP) isolated from influenza virus. Scale bars are panel B (10 nm) and panels C and D (5 nm). (E) Schedule for injection of mice with purified rNP and viral NP. Female BALB/c mice were used. Blood was drawn on days 0, 14 and 35. Aliquots of sera at terminal bleed day 35 were pooled for immunoassays. The route of injection was intramuscular (IM) at days 0 and 21. A total of 100 micrograms of NP were given to each mouse. (F) Binding activity of mouse sera to rNP following immunization with PBS (black), rNP (blue) or viral NP(red) assessed by ELISA. (G,H) Probing the specificity and reactivity of anti-rNP (panel G) and anti-viral NP (panel H) mice sera to viral lysates from purified influenza viruses that were denatured and subjected to SDS-PAGE before transfer to nitrocellulose membranes. Samples were probed with anti-rNP and anti-viral NP polyclonal mouse sera as primary antibody. Immunoblot standard (std) is shown along with influenza viral samples that were A/Puerto Rico/8/34 (H1N1), A/Victoria/3/75 (H3N2), and B/Lee/40. Molecular weight standards (std) and lysozyme negative-control are denoted. (I) Immunoblot using anti-rNP mouse sera to detect decreasing concentrations of rNP protein.

Pooled sera from day 35 was used in ELISA as primary antibody to test the reactivity and specificity of the anti-rNP and the anti-viral NP sera to rNP (Fig. 2F) and viral lysates (Fig. G, H). Mouse sera raised to either viral NP or to rNP both could detect rNP by ELISA (Fig. 2F), and there appeared to no statical difference in the immunogenicity to rNP versus viral NP (Fig. 2F. Fig. S1C). Immunoblots probed with either anti-rNP or anti-viral NP mouse sera indicated NP bands at 55 kDa (Fig. 2G, 2H). However, when probing reactivity of anti-NP sera against viral lysates via immunoblotting, the anti-rNP mouse sera appeared to be more specific to H3N2 (Fig. 2G). In contrast, the anti-viral RNP sera (viral NP) appeared to elicit more cross-reactivity to H1N1, H3N2, influenza B viral lysates (Fig. 2H). This may be because the rNP is purified with no other viral proteins while viral NP contains NP as the major component with trace amounts of other viral protein such as HA. In addition, further analysis of the anti-rNP mouse sera indicated that some of the antibodies appear to bind linear epitopes in that rNP bands were detected in an immunoblot format in which rNP is denatured (Fig. 2I). These results suggested that an appropriate NP antibody could be used to quantitate NP in vaccine formulations.

### 2.3. Analysis of purified anti-NP mouse monoclonal antibody HB65

We selected mAb HB65 as a representative NP-targeted antibody because it has been investigated by several groups for diverse applications, and it is available commercially. We obtained HB65 antibody by purification from HB65 hybridoma cell culture supernatant by purification using protein G affinity.

To determine the purity and disulfide bonding status of our purified anti-NP mouse monoclonal antibody HB65, we used SDS-PAGE analysis under reducing and non-reducing conditions. Purity was confirmed by observing the number of protein bands and the relative portions of heavy and light chain bands at expected molecular weight migrations as compared to standards. Under reducing conditions bands were observed at apparent molecular weights of 20 kDa, and 50 kDa which are the approximate molecular weights for IgG antibody light chains and heavy chains, respectively (Fig. 3A). Antibody HB65 was judged to be greater than 95% pure by densitometry. Assessment of protein purity and homogeneity was important before further analysis on mAb HB65, such as by electron microscopy and the use of mAb HB65 in immunoassays.

**Fig. 3.**
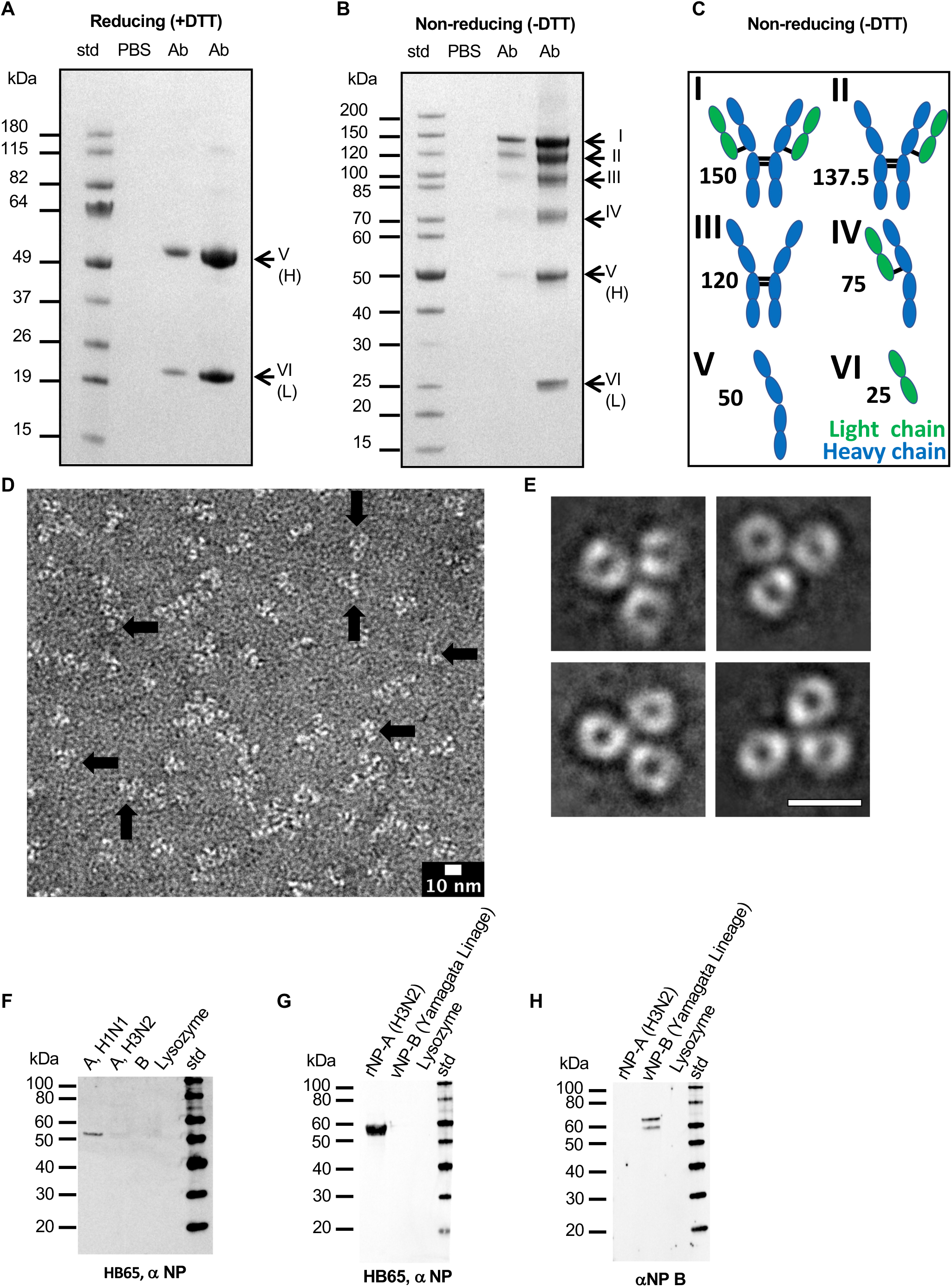
Analysis of the purity, structure, and reactivity of anti-NP mouse monoclonal antibody HB65. (A) SDS-PAGE analysis of mAb HB65 (Ab) under reducing conditions (+dithiothreitol (DTT)) and (B) under non-reducing conditions (-DTT). Samples are molecular weight standards (std), PBS, and different concentrations of antibody (Ab). In panel A, arrows point to major bands ascribed to reduced and separated heavy and light chains. In panel B, arrows point to both heavy and light chains proteins ascribed as having combinations of formed and non-formed inter-chain disulfide bonds based on apparent molecular weights comparison of IgG with constituent heavy and light chain components (I-VI). (C) Schematic depicting possible molecular components of an IgG molecule based on variable inter-chain disulfide bond formation between heavy and light chains. Heavy chains are in blue and light chains are in green. Each component immunoglobulin (Ig) subunit is represented by an oval. Black lines represent inter-chain disulfide bonds. Species depict (I) IgG with all inter-chain disulfide formed, (II) two heavy chains with one light chain, (III) Two heavy chains, (IV) one heavy and one light chain, (V) one heavy chain, and (VI) on light chain. The approximate molecular weight of each species is denoted and based on a single IgG domain fold of about 12.5 kDa in molecular weight. This results in estimated weights of 150 kDa, 137.6 kDa, 120 kDa, 75 kDa, 50 kDa, and 25 kDa for components I-VI, respectively. (D) Electron microscopy image with observed molecular complexes stained with 0.5% Uranyl Acetate. Arrows point to some isolated antibodies. (E) Example images of 2D class-averages of IgG HB65. For panels D and E contrast is with protein in white with scale bars, 10 nm. (F,G). Immunoblots to probe for the detection of NP by mAb HB65 in different samples such as viral lysates (H1N1, H3N2, B/Lee) (panel F) and rNPs (rNP-A, rNP-B) (panel G). (H) Commercial anti-influenza B NP sera was a detection control for rNP-B.

Under non-reducing conditions a ladder of bands was observed for mAb HB65. An IgG has a molecular weight of about 150 kDa. A band was observed at this size (Fig. 3B, top band I). However, under non-reducing conditions other bands appeared below the 150 kDa band (Fig. 3B bands II-VI). Six hypothetical schematic models were constructed, and their estimated molecular weights were based on general IgG structure. Heavy and light chain species were given different levels of inter-chain disulfide bond formation to produce six schematics (Fig. 3C). Model schematics of the different protein species were constructed by assigning an IgG domain fold with a molecular weight of 12.5 kDa. Thus, the six schematics could correspond to the six observed bands for mAb HB65 under non-reducing conditions (Fig. 3B vs 3C, species I-VI). Based on densitometry the relative amounts of the species were in a mostly decreasing order: (26%, band I, 150 kDa), (20%, band II, 137.5 kDa), (16%, band III, 120 kDa), (12%, band IV, 75 kDa), (15%, band V, 50 kDa), (11%, band VI, 25 kDa). MAb HB65 had been reported to be an IgG2a. Edman sequencing of the N-terminal of the light-chain was used as direct chemical analysis of the sample to confirm it as an antibody sequence. A sequence of twelve amino acid residues of the N-terminal protein sequence was NIVMTQSPKSMS. A subsequent protein Basic Local Alignment Search Tool (BLAST) database search identified a series of homologous sequences that belonged to mouse light chain antibodies. Hits with 100% sequence identity to the HB65 peptide were immunoglobulin kappa light chain (Mus musculus) accession #CAA24192.1, Antibody G196 accession #5H2B-B, and anti-Flavivirus Fab4g2 accession #1UYW-L.

### 2.4. Electron microscopy of anti-NP mAb HB65 and image analysis

To address the question of what effects would the observed differential disulfide bonding pattern have on HB65 IgG structure, we analyzed the structural integrity and organization of mAb HB65 by 2D-electron microscopy and observed predominantly monodisperse molecular complexes, yet some complexes were found overlapping (Fig. 3D). The complexes appeared as multi-lobed structures with some instances of presenting Y-shaped molecules (Fig. 3D), consistent with classical IgG molecular organization. To arrive at a representative average image of HB65 molecules, 2D class average image analysis was pursued. Hundreds of individual complexes of mAb HB65 molecules were picked (N=561) and subjected to reference-free 2D class averaging. Class averages yield images of higher contrast and detail than raw images alone (Fig. 3D vs. 3E). The 2D classes displayed images of Y-shaped antibody molecules with three arms (Fig. 3D). At the resolution of negative-stain EM, we cannot differentiate between Fab and Fc arms, yet still the characteristic Y-shape molecular organization of Ig domains is clearly observed. From these results we confirm that purified anti-NP mAb HB65 is an intact IgG Y-shaped molecule despite not all disulfide bonds being formed.

### 2.5. Probing for anti-NP mAb HB65 specificity with influenza type A and type B viral lysates and recombinant NP

To characterize the interaction and specificity of anti-NP mAb HB65, immunoblot and ELISA were carried out using influenza viral lysates, rNP, and M1 proteins. Immunoblots indicated that the epitope for mAb HB65 appeared to be linear and specific for NP of type A influenza viruses. The strongest detections of NP bands by mAb HB65 via immunoblotting were for type H1N1 viral lysate (Fig. 3F) and H3N2 recombinant NP (Fig. 3G). The reactivity of HB65 to NP appeared specific in that HB65 was not reactive to a lysozyme negative control (Fig. 3G, 3H). Recombinant NP from influenza B virus was not detected by HB65 (Fig. 3G). We compared the primary amino acid sequence of NP from influenza A and B to explain the lack of binding and found the sequence identity was 34.5% and the sequence similarity was 51.6% (Fig. S2). Indicative of the specificity of HB65 via immunoblot of H1N1 viral lysate, mAb HB65 did not detect to comparable levels other viral structural proteins like matrix protein M1 (∼30 kDa expected) or HA (∼70 kDa expected) (Fig. 3F).

To explore if the HB65 epitope required eukaryotic post-translational modification, we used rNP isolated from bacteria to probe for reactivity to mAb HB65 via immunoblot (Fig. 4A). MAb HB65 could detect bacterially expressed NP under both reducing and non-reducing conditions (Fig. 4A). The specificity of mAb HB65 of NP was further confirmed by additional analysis using recombinant matrix M1 protein in ELISA (Fig. 4B) and HA containing commercial influenza vaccines in ELISA and immunoblotting (Fig. 4C, 4D) (Fig. S3C vs S3D). MAb HB65 was specific of NP and did not cross-react with matrix M1 in ELISA (Fig. 4B). Monoclonal antibody HB64 is an anti-M1 antibody, and it too was specific for M1 and did not cross react with NP in ELISA (Fig. 4B).

**Fig. 4.**
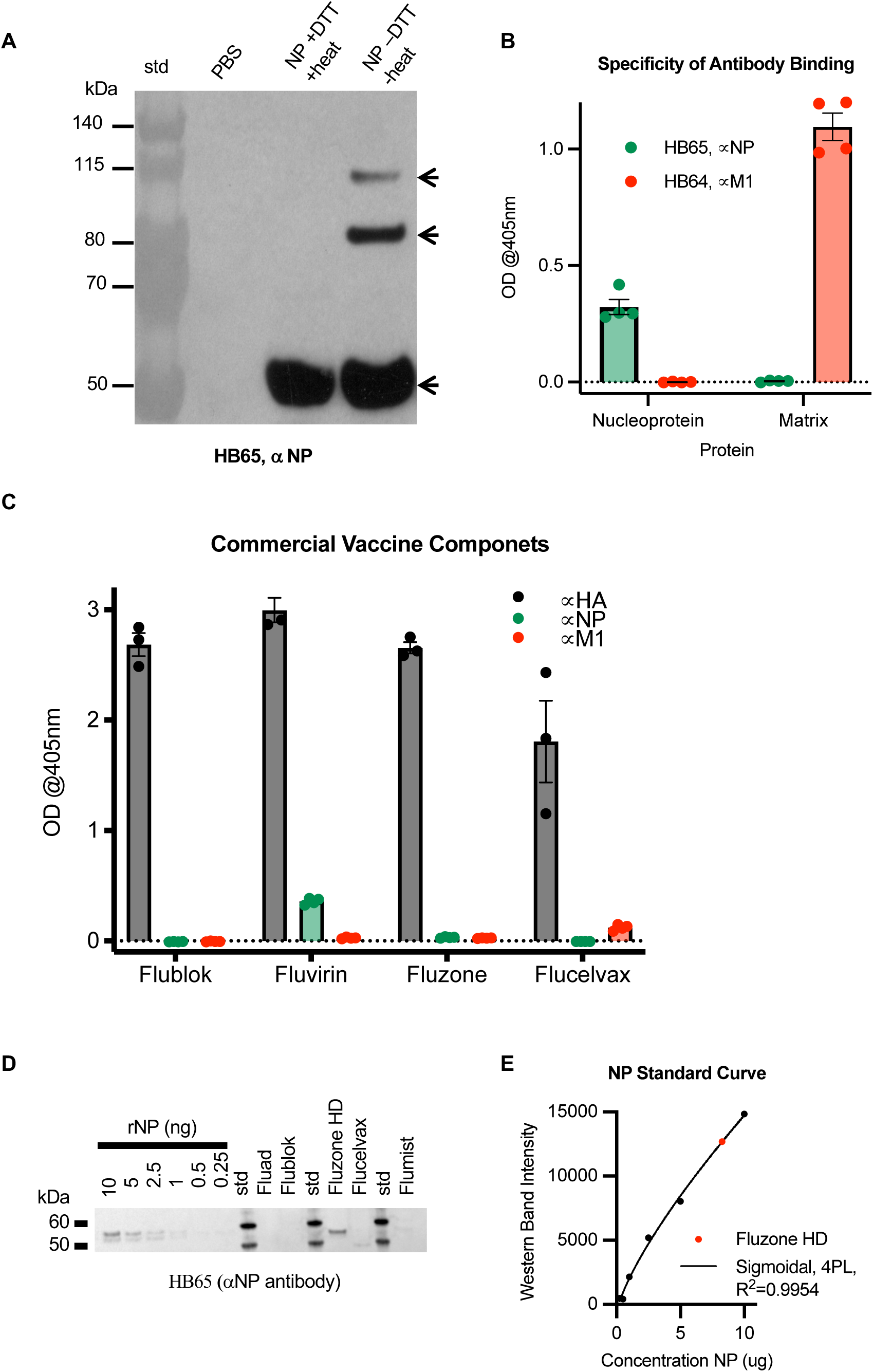
Probing the reactivity and specificity of anti-NP mAb HB65 to detect and quantitate NP in commercial influenza vaccines. (A) Immunoblot analysis of mAb HB65 reactivity to NP under reducing (+DTT) and denaturing (+heating) conditions compared to NP non-reducing (-DTT) and no heating before SDS-PAGE and sequent transfer to nitrocellulose membrane for immunoblotting. Recombinant NP (A/Puerto Rico/9/1834 (H1N1)) was purified from bacteria. Arrows indicated a ladder of NP bands detected by mAb HB65 in the NP -DTT and NP -heat sample. (B) ELISA probing for the reactivity and specificity of mAb HB65 (green) with rNP and Matrix (M1) proteins (baculovirus expression, H3N2 virus) versus an anti-M1 mouse monoclonal antibody (HB64) (red) as control. (C) Probing the use of mAb HB65 to detect NP (green) in commercial influenza vaccines and compared to anti-H1 antibody (gray) and anti-M1 antibody HB64 (red). Commercial influenza vaccines were Flublok, Fluvirin, Fluzone HD, and Flucelvax. (D) Immunoblot with deceasing amounts of rNP and with different commercial influenza vaccines. Standards are denoted (kDa). (E) NP standard curve based on immunoblot band intensity of NP in panel D.

Because mAb HB65 (anti-NP) and mAb HB64 (anti-M1) were found specific, their potential uses as diagnostic antibodies were tested for the detection of NP and M1 in commercial influenza vaccines. Four commercial influenza vaccines were tested: Flublok, Fluvirin, Fluzone, Flucelvax. By ELISA NP was detected in Fluvirin, and M1 was detected in Flucelvax (Fig. 3C). As expected, the strongest level of detection was observed with a polyclonal anti-HA antibody (H1), since HA is the major immunogen in theses vaccines (Fig. 4C). Since anti-NP mAb HB65 has a linear epitope detectable by immunoblot, HB65 was used to quantitate the relative amounts of NP in commercial vaccines (Fig. 4D, 4E) (Fig. S3). Based on a dilution series of NP, at least 1 nanogram of NP could be detected by anti-NP mAb HB65 (Fig. 4, Fig. S3), Among the commercial influenza vaccines tested via immunoblot, NP was detected in abundance only in Fluzone high dose (HD) (Fig. 4D). Based on a NP standard curve (Fig. 4E) Fluzone HD has 3.25 μg of NP/240 μg HA (Fig. 4D, S3). These results confirm anti-NP mAb can detect NP in commercial influenza vaccines in both ELISA and immunoblot immunoassay formats (Fig. 4C, 4D).

The ability of mAb HB65 to detect NP in complex mixtures may be due to its high affinity. Bio-layer interferometry (BLI) was used to measure the binding interaction between mAb HB65 and rNP. The interaction was dose-dependent in that increased concentrations of rNP increased the level of binding by immobilized mAb HB65 (Fig. S4). The mAb HB65 and rNP interaction had a measured K_D_ of 120 ± 21 nM (Fig. S4). Although mAb HB65 did not protect naïve mice from H1N1 influenza infection via intraperitoneally passive transfer experiments when compared to anti-HA polyclonal antibody (Fig. S5), these results suggests that combined mixtures of polyclonal anti-NP and anti-HA antibodies might provide improved protection in future studies of anti-NP antibodies.

### 2.6. Anti-NP mAb HB65 epitope mapping via deletion library and molecular modeling

To further understand the epitope determinant of mAb HB65, deletion mapping was used to map the general region of the mAb HB65 epitope within the NP sequence. This strategy was used because the mAb HB65 bound to denatured NP via immunoblot, strongly suggesting a linear epitope (Fig. 4D). Six constructs including wild-type NP and five deletions were engineered by sequentially deleting segments of 100 non-overlapping amino acids from the NP primary sequence (Fig. 5A). Because mAb HB65 bound recombinant NP expressed in bacteria, deletion constructs were also expressed in bacteria. NP was purified by nickel-affinity chromatography via an N-terminal histidine tag. NP deletion constructs were named according to the amino acids deleted: WT 1-498, Δ1-100, Δ101-200, Δ201-300, Δ301-400, and Δ401-498. The deletions represented spanned both helical and loop structural elements when mapped onto the monomer structure of NP (Fig. 5C). In immunoblot analysis using the NP proteins, mAb HB65 detected full-length NP WT 1-498 and deletion constructs in the C-terminus, Δ301-400, and Δ401-498 (Fig. 5D). N-terminal deletion constructs, Δ1-100, Δ101-200, Δ201-300 were not detected by HB65 (Fig. 5D). Thus, the epitope for mAb HB65 was consistent with mapping to the N-terminal half of the NP sequence (Fig. 5D).

**Fig. 5.**
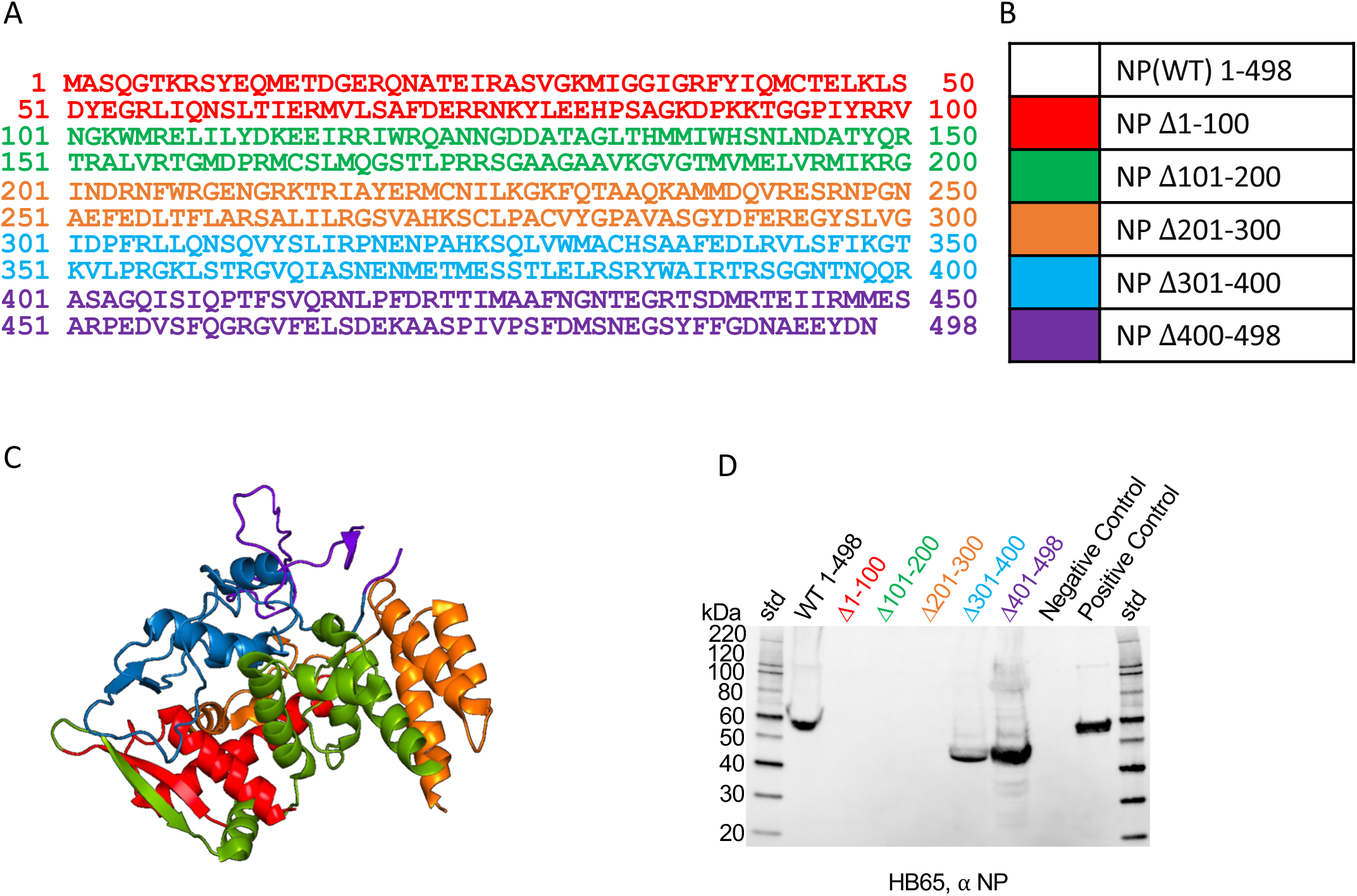
Epitope mapping of anti-NP mAb HB65 using a recombinant NP deletion library. (A) Sequence for influenza NP (A/California/07/2009 (H1N1)). Starting from the N-terminus, blocks are color coded in groups of 100 residues until the last block which contains only 98 residues at the C-terminus. Corresponding residues missing for the crystal structure of NP (PDB 2IQH) are underlined. (B) Color coded key identifying the deletion construct and region deleted. (C) Structure of NP monomer (PDB 2IQH) shown as a ribbon diagram with corresponding-colored regions marked for deletion. N- and C-termini are labeled. (D) Immunoblot of rNP deletion library proteins purified from bacteria and probed with mAb HB65. Standards are denoted (std). Controls are positive control (rNP) and negative-control (lysozyme).

### 2.7. Anti-NP mAb HB65 sequence and molecular modeling

To further understand the sequence and organization of mAb HB65, a homology model was constructed, starting from confirmation of the DNA sequence. RNA isolated from hybridoma cells was used to produce cDNA sequences of the variable light (VL) and variable heavy (VH) domains (Fig. 6A, 6C). Corresponding protein sequence information was derived by computationally translating the DNA into corresponding protein sequences (Fig. 6B, 6D). The protein sequences were used in homology modeling to derive a model structure for the VH and VL IgG domains of mAb HB65 (Fig. 6E). Three different regions were identified from the sequences: leader sequences, framework regions, and the intervening complementarity determining regions (CDRs) for light and heavy chain regions (Fig. 6). The heavy chain sequence of HB65 was found to be unique upon BLAST query of GenBank. The complementary determining region 2 (CRD2) of the heavy chain was among the longest among CDRs (Fig, 6D). The sequence of the VH domain of mAb HB65 was found to a match from BLAST analysis (80.2 % identity) corresponded to mouse antibody D12, which binds to a region of the MERS coronavirus spike glycoprotein (Wang et al., 2015). Among the antibody sequences that mAb HB65 shared similarities with, their respective antigens did not share similarity, suggesting that the HB65 germ-line sequence is not necessarily predisposed or biased to recognize NP antigen.

**Fig. 6.**
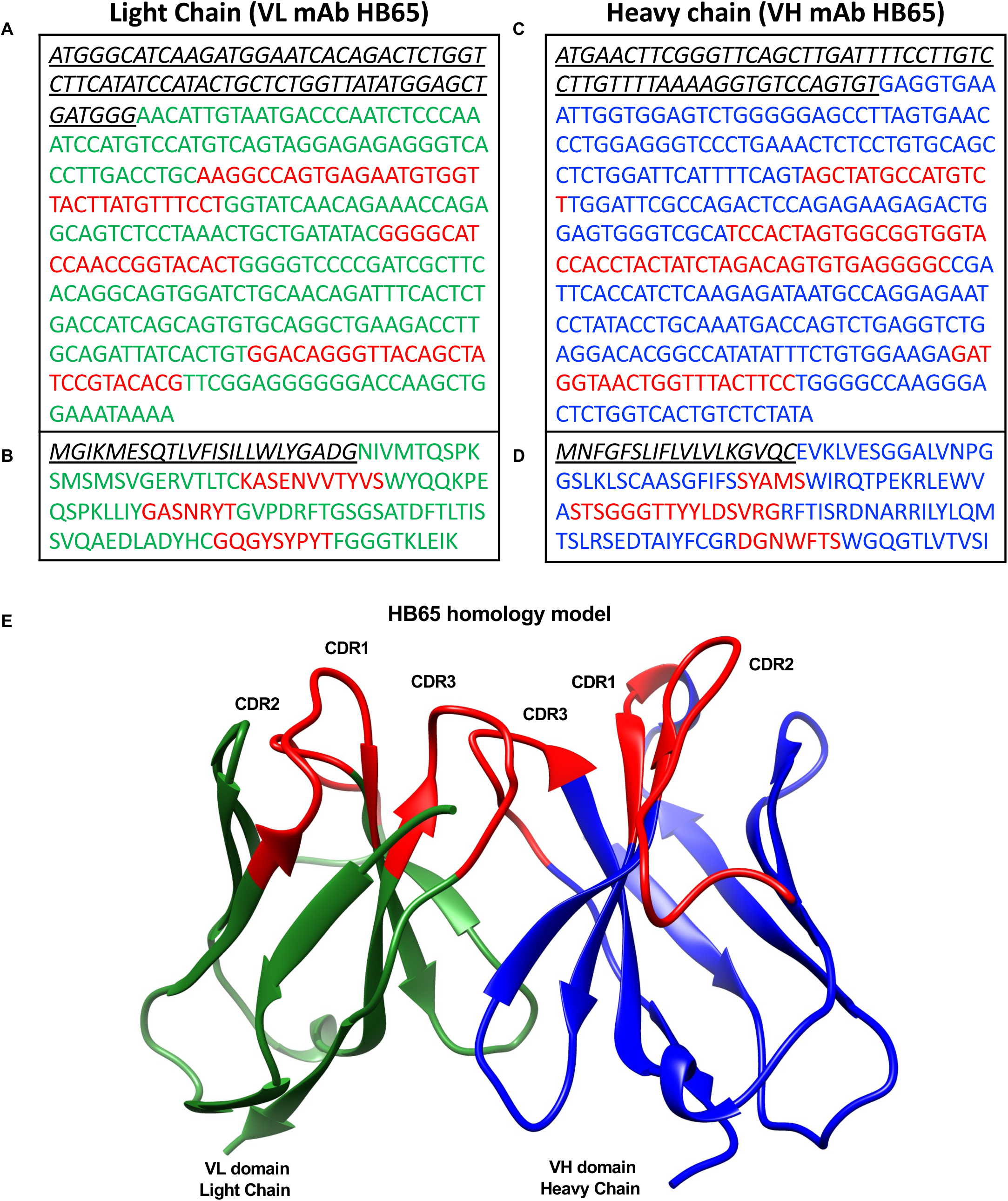
Sequences of variable light and heavy regions of anti-NP mAb HB65 and homology modeling of structure. (A) DNA sequence of mAb HB65 variable light chain (VL domain) and (B) corresponding derived protein sequence. Leader sequence is in black with VL in green and the complementarity-determining regions (CDRs) in red. (C) DNA sequence of mAb HB65 heavy light chain (VH domain) and (D) corresponding derived protein sequence. Leader sequence is in black with VH in blue and the complementarity-determining regions (CDRs) in red. (E) Homology model of mAb HB64 variable domains for the light and heavy chains. The VL domain is green with the VH domain in blue and CDRs (CDR1, CDR2, CDR3) for each domain are colored red.

## 3. DISCUSSION

There is increasing evidence that internal viral proteins can elicit antibodies that can be used as important diagnostic tools. Influenza nucleocapsid protein (NP) is an example of such an internal antigen, as NP is the major protein component of packaged genomic viral RNPs in the center of the virion. And yet there have also been reports of NP on the surface of virus infected cells and NP eliciting antibodies which facilitate antibody-mediated immune responses and protect from lethal infection (Bodewes et al., 2013; Carragher et al., 2008; LaMere et al., 2011; Vanderven et al., 2016; Yewdell et al., 1981). The ability of NP to elicit antibodies makes NP of interest for both the development of diagnostic assays and the development of more efficacious influenza vaccines. While NP is highly conserved, it is unknown how different preparations or presentations of NP antigens could affect NP immunogenicity, NP epitope display, and the diagnostic abilities of both NP antigens and elicited anti-NP antibodies.

As a surrogate for NP preparations in the context of viral infection, we first examined serum from influenza-infected mammals to determine if different influenza type A strains could elicit detectable levels of NP antibody. We found that H5N1 convalescence human sera had antibodies to NP, and that some of these epitopes were linear (Fig. 1A, 1B). Anti-NP antibody levels were not significantly different between the H5N1 sera groups, although they followed the trend of HAI titer levels (Fig. 1A).

Additionally, we detected the presence of anti-NP antibodies in sera from groups of ferrets infected with H1N1, H3N2 and H7N9 viruses, which represent influenza A viruses from different subtypes representing group 1 (e.g. H1) and group 2 (e.g. H3, H7). All infected ferret sera recognized linear epitopes on NP, as indicated by immunoblot analysis with denatured NP antigen (Fig. 1F, 1G, 1H). The ferret sera from H3 infection showed the strongest binding to NP as judged by ELISA (Fig. 1E), but we also note that the NP detection antigen sequence was derived from a H3N2 virus and thus was expected to be stronger than cross-reactive binding. In terms of using NP as a diagnostic antigen, we find cross-reactivity of anti-NP antibodies, but our work also indicates that strongest diagnostic responses would be obtained utilizing a NP antigen reagent comprised of a mixture of NP antigens from a selection of viral subtypes, such as H1, H3 and H7.

In addition to subtype differences, we explored if the source of the NP antigen, either recombinant NP (rNP) or viral NP (vNP), may also contribute to differences in the detection of NP antigen and anti-NP antibodies. Previous studies with hemagglutinin (HA)-based vaccines indicated that the structural display of HA is variable between vaccines, and that HA complex organization can modulate the elicitation of cross-reactive antibodies (Myers et al., 2023). We asked if NP antigen organization affected NP detection and anti-NP antibodies, speculating that the molecular organization of NP may modulate which NP epitopes are displayed, thus affecting immunogenicity and detection efficacy by anti-NP antibodies. For example, there are reports of variability and therefore inconclusive results from rapid influenza diagnostic tests (RIDTs) that detect influenza viral NP antigen (Balish et al., 2013; Bose et al., 2014). RIDTs are relied upon both for surveillance of influenza animal viruses and for point-of-care with patients in clinical practices. Many RIDTS utilize the principle of lateral flow immunoassays, where the patient sample flows along a nitrocellulose membrane in a cassette designed with bands of conjugated labels and antibodies for detection via a visual readout of appearing bands. For influenza virus RIDTS assays, anti-NP antibodies or NP protein are often selected as the targets fixed to the nitrocellulose membrane.

Similar to influenza, diagnostics for other prevalent viruses such as SARS-2 (COVID-19) use recombinant SARS nucleocapsid protein (N) and anti-N monoclonal and polyclonal antibodies for band visualizations (Burbelo et al., 2020; Dugan et al., 2021; Pollock et al., 2021). Moreover, anti-NP antibodies have been reported to play a role in influenza immunity (Berthoud et al., 2011; Kaminski and Lee, 2011; Vanderven et al., 2016) and even perhaps autoimmunity in narcolepsy (Ahmed and Steinman, 2017). However, characterization of monoclonal antibodies to NP is underdeveloped compared to antibodies to hemagglutinins (Burton et al., 2012; Caton et al., 1982; Gerhard et al., 1981; Joyce et al., 2016).

We utilized the anti-NP mouse mAb HB65 for our experiments because HB65 was one of the first reported antibodies to NP (Yewdell et al., 1981) and has been reported to have a wide range of applications, including flow cytometry and immunofluorescence (Yewdell et al., 1981). We characterized the purified mAb HB65 (Fig. 3) and its interaction with NP (Fig. S4), then quantified NP in various samples including commercial influenza virus vaccines (Fig. 4). Consistent with the reactivity of naturally infected mammal serum, binding of mAb HB65 to NP appeared to be mediated by a linear epitope (Fig. 4D). Our results showing mAb HB65 reactivity to NPs from different influenza type A virus subtypes (Fig. 3F, 3G) could explain why in previously reported work mAb HB65 was able to detect different viral subtypes by immunohistochemistry methods (Nicholls et al., 2012).

Our analyses with anti-NP mAb HB65 have several implications for NP-based diagnostic tests. Our immunoblotting procedure mimics that of RIDT as we use a lateral flow technique to apply antibodies to a nitrocellulose membrane displaying target protein. Our results indicated that mAb HB65 can be utilized in a lateral flow technique to detect recombinant NP as well as NP from lysed influenza virus, which we have referred to as viral NP (Fig. 3F, 3G). The ability of anti-NP polyclonal antibodies and mAb HB65 to detect NP in different structural states such as linear polypeptides, NP rings and NP filaments (Fig. 2, Fig. 3, Fig. 4) enables these antibodies to be used for additional diagnostic tests, such as the quantification of NP in commercial influenza vaccines (Fig. 4C-4E). Next generation commercial vaccines are including NP by design and are finding improved results (Lo et al., 2021; Pardi et al., 2022; Sayedahmed et al., 2024). Thus, future standardization of NP concentrations in commercial influenzas vaccines, such as we have shown with mAb HB65, may become an important priority. We have observed that HB65-like antibodies are useful for quantitating NP in a variety of formats, including in a denatured state such as immunoblot (Fig. S3), in an absorbed state such as in ELISA (Fig. 4C), or in a soluble state with native tertiary and oligomeric structure such as in biolayer interferometry (BLI) format (Fig. S4).

Following from previous applications of BLI to other systems (Cate et al., 2021; Zhang et al., 2021), our results could aid in the automation and processing of both clinical and surveillance samples leading to further development of NP-based diagnostics assays for influenza viruses.

Diagnostics probes are now being engineered based on the sequences of lead antibodies. Nanobodies and biotinylated detection reagents are being engineered starting from a pre-existing natural antibody (Du et al., 2019; Ji et al., 2022). Foundational work on anti-NP sera and anti-NP monoclonal antibody sequences is not as well developed as equivalent work on anti-HA sera and anti-HA monoclonal antibody sequences. Thus, to provide further support for the use of a HB65-like antibody as a diagnostic, we have characterized the epitope footprint for mAb HB65 (Fig. 5) and determined the sequence of mAb HB65 (Fig. 6). The mAb HB65 epitope maps to the N-terminal of region of NP (Met1-Gly300) based on protein deletion constructs of NP (Fig. 5). In future studies, it will be interesting to use the sequence of mAb HB65 to engineer nanobodies and biotinylated antigen as done with other antibody sequences (Du et al., 2019; Ji et al., 2022). Also, in future studies, it will be of great value to identify other monoclonal antibodies to NP, especially those which may have non-overlapping NP epitopes. These antibodies could provide anti-NP capture or detection in tandem with HB65-like antibodies, thus aiding in the development of more robust assays to detect influenza virus infection.

## 4. CONCLUSION

In conclusion, our results indicate that NP is immunogenic in mammals and that NP elicits anti-NP antibodies. Our work suggests that anti-NP sera are cross-reactive across most strains within an influenza type, but binding can be weaker for more distantly related influenza subtypes. Mixtures of NPs or anti-NP antibodies derived from just a few influenza type A subtypes are likely to span all sequences necessary for diagnostic efficacy. We further characterized the classical anti-NP mouse monoclonal antibody mAb HB65, which is used in the influenza research field as a research reagent but lacks extensive characterization as a diagnostic antibody reagent. We observed that mAb HB65 had a linear NP epitope specific for type A influenza viruses, which it bound with high affinity (120 nM). Furthermore, mAb HB65 could detect NP in some commercial influenza vaccines. Our results create the expectation that further work characterizing the concentration of NP within vaccine preparations will be of great interest as more vaccines seek to include NP due to its conserved sequence compared to other influenza proteins like antigenically variable hemagglutinin (HA).

## 5. MATERIALS AND METHODS

### 5.1. Viruses and sera

Influenza viruses used for SDS-PAGE and then subsequent immunoblots (western blots) were purified influenza viruses: A/Puerto Rico/9/34 (H1N1), A/Victoria/3/75 (H3N2), and B/LEE/40) (Charles River Laboratories, INC., North Franklin, CT). Viruses were propagated in and purified from 10- to 11 days old embryonated hen’s eggs as previously reported (Harris et al., 2013). Sera from influenza infected ferrets was kindly provided by the International Reagent Resource (IRR). Three different ferret antisera were to influenza viruses: A/California/07/2009 (H1N1) pdm09, A/Rhode Island/01/2010 (H3N2), A/Anhui/1/2013 (H7N9). H5N1 polyclonal antisera were human reference antisera to influenza virus A/Indonesia/5/2005 (H5N1) (Biodefense and Emerging Infections Research Resources Repository (BEI Resources).

### 5.2. Mouse immunogenicity and sera and passive-transfer

All animal experiments were performed under protocols approved by the Animal Care and Use Committees of the National Institute of Allergy and Infectious Diseases. The recombinant NP (rNP) used as a mouse immunogen was from a baculovirus expression system (IRR), This NP had a N-terminal histidine tag, and the NP sequence was from Influenza A/Brisbane/10/2007 (H3N2) (IRR). This Influenza NP immunogen was judged to be more than 98% percent pure by SDS-PAGE. Mice were female BALC/c mice (Charles River Laboratories) aged 8-10 weeks. Two groups of mice (n=5) were anesthetized by isoflurane inhalation. One group was a PBS control group, and the other group was a NP injection group. NP was in sterile PBS at a concentration of 0.5mg/ml and mice were injected by intramuscular (IM) route with 50 μl per each hind leg on days 0 and 21 resulting in a total of 200μl (100μg) of NP injected into each mouse. Sera was collected on day 0 and 14 by tail vein bleeds, and on day 35 by terminal bleeds.

Supplemental passive transfer experiments in mice using anti-NP mouse monoclonal antibody HB65 were like those previously described using anti-HA monoclonal antibodies (Myers et al., 2023). Briefly, passive-transfer of 400 μl (350 μg/mouse) of anti-NP mAb HB65 (0.875mg/ml) or 400 μl of anti-H1 HA vaccine (Flublok) sera were administered intraperitoneally (IP) 18 hours before intranasally challenged with influenza virus (A/Michigan/45/2015) (H1N1).

### 5.3. Purification of recombinant NP from bacteria and viral NP from influenza virus

For production and purification of NP from bacteria, genes encoding nucleoproteins (NP) for influenza A/PR/8/34 (H1N1) (type A NP) and B/Lee/40 (type B NP) in a pCMV-Entry vector (OriGene Technologies, Rockville, MD), were individually subcloned using AscI and Mlu I restriction sites into the bacterial expression vector pEx-N-Hi. This expression vector codes for a N-terminal histidine tag.

Resulting plasmids were transformed to *E. coli* expression cells Rosetta 2 (DE3) cell for protein expression. For each type A NP and type B NP transformed colonies were added to 1 liter LB Broth with appropriate antibiotics and at an optical density of 0.6 at 600 nm protein expression was induced the addition of a final concentration 1 mM IPTG. Subsequently after 3.5 hours, cells were pelleted from culture medium by centrifugation (2000 rpm at 14 °C for 20 minutes with a SX4750A rotor). Pellets were resuspended in ice-cold PBS with protease inhibitor cocktail and disrupted by sonication to produce crude lysates. Supernatants containing NP were obtained from lysates by centrifuged at 12,000 x g for 20 minutes at 4 °C. Type A and type B nucleoproteins were purified from their respective supernatants by affinity chromatography (immobilized metal affinity chromatography) using gravity-flow columns (His GraviTrap TALON) with bound protein eluted with increasing concentration of imidazole. Fractions containing purified NP as assessed an SDS PAGE were concentrated using a centrifugal filter device (Amicon Ultra-4) coupled with buffer exchange using PBS. NP samples were stored at 4 °C at 0.8mg/ml until further use in immunoblot and ELISA experiments. Isolation of viral NP (i.e. filamentous genomic RNPs) from influenza virus has been described previously using detergent lysis and sucrose gradients (Gallagher et al., 2017).

### 5.4. Immunoassays (ELISA)

For ELISA experiments, indirect ELISA was performed. Antigen samples to be probed were rNP, influenza matrix protein (M1) and some commercial influenza vaccines. Commercial influenza vaccines were obtained from their respective companies/distributors and consisted of 2016 vaccines: Fluzone, Flublok, Fluvirin, and Flucelvax. For 2017 influenza vaccines the samples were Flublok, Flublok Quadrivalent, Fluzone High Dose (HD), Flucelvax, and Fluad. Primary antibody included pooled sera of ferrets and humans infected with different subtypes of influenza viruses, mouse sera from mice immunized with recombinant NP and purified mouse monoclonal antibody HB65 (mAb HB65). Primary antibodies were diluted 1:1000 for analyses and secondary according to manufacturer’s instructions. For ELISA, antigens were coated on 96-well microplates and incubated at 4 °C for 12 hours and then blocked with 5% non-fat dry milk in 1X PBS at 37 °C for 2 hours. The plates were washed 2 times with 1X PBS and then incubated with primary antibody at 4 °C for 12 hours. The plates were washed 4 times with 1XPBS and then incubated with the appropriate secondary antibody conjugated to horseradish peroxidase at 37 °C for 2 hours. The plates were washed four times with 1X PBS and then color was developed with 1-Step ABTS (Thermoscientific, Rockford, IL). The absorbance was measured with a plate reader at 405 nm. ELISA assays were performed in quadruplicate.

### 5.5. Immunoassays

For immunoblots (western blots), samples were separated by SDS-PAGE before transfer to nitrocellulose membranes. Samples with antigens to be probed were influenza viruses, rNP and M1. Also, samples for immunoblot analysis included some commercial influenza vaccines. Primary antibody included pooled sera of ferrets and humans infected with different subtypes of influenza viruses. Primary antibody also included mouse sera from mice immunized with rNP and purified mAb HB65. Secondary antibodies were goat anti-ferret IgG, goat anti-human IgG, and goat anti-mouse IgG that were fluorescent-labeled secondary antibodies. After SDS-PAGE, respective samples were transferred to nitrocellulose membranes via the iblot transfer system (Thermofisher) and probed with primary antibody and florescent tagged secondary antibody via lateral flow (ibind system, Thermofisher). Membranes were visualized on the c600 imager (Azure biosystems).

### 5.6. Purification of anti-NP monoclonal antibody HB65

For purification of mAb HB65, deposited hydridoma cells for mAb HB65 were acquired from the American Type Culture Collection (Manassas, VA). Hydridoma cells were grown by standard procedures. In brief, hybridomas were subcultured in DMEM with 10% FBS. Antibody in culture supernatants from hybridoma cells were purified by standard affinity chromatography techniques. Briefly, each sample was diluted in column equilibration/wash buffer (10 mM NaPO4, 150 mM NaCl pH 7.0). The antibodies were purified on Protein G columns and eluted with 100 mM glycine pH 2.5. Purified antibody fractions were neutralized to pH 7.4 with 4.0 M Tris pH 8.0 and stored at 4 °C at 3.5mg/ml until further use. Purity of antibody was assessed by SDS-PAGE. Chemical identification and confirmation of the sample as an antibody was assessed by N-terminal protein sequencing by Edman sequencing of the putative light chain protein band.

### 5.6. Sequence information and molecular modeling for anti-NP monoclonal antibody **HB65**

To obtain additional sequence information for mAb HB65, hybridoma cell pellets were used to extract RNA for cDNA synthesis with subsequent sequencing of the cDNA for genes for variable domains of the heavy and light chains of mAb HB65 (GenScript, Piscataway, NJ). Briefly, total RNA was isolated from the mAb HB65 hybridoma cell pellets and reverse transcribed into cDNA using primers for heavy and light chains. Amplified antibody fragments were separately cloned into a standard cloning vector, colony PCR screening was performed to identify clones with inserts of correct sizes. Individual positive clones with correct variable heavy (VH) and variable-light (VL) insert sizes were sequenced. The Basic Local Alignment Search Tool (BLAST) was used to search for similar sequences within GenBank to VL and VH domains of mAb HB65. Molecular modeling of mAb HB65 was carried out using the Swiss-model server (Biasini et al., 2014). The model was for the variable light (VL) and variable heavy (VH) domains. First, protein sequences were computational derived from the determined cDNA sequences for the VL and VH domains. These sequences were submitted to the Swiss-model server. The coordinate models for mAb HB65 VH and VL domains were two Ig domains forming the anti-NP paratope. For sequence alignments EMBOSS lite and ClustalW2 were used with Chimera for coordinate and sequence visualization (Pettersen et al., 2004).

### 5.7. Bio-layer interferometry of NP and anti-NP monoclonal antibody HB65

The interaction between the mouse anti-NP monoclonal antibody HB65 and rNP was determined by bio-layer interferometry (BLI) using an Octet Red 96 instrument according to manufacturer’s suggested protocol (Sartorius (ForteBio Inc., Menlo Park, CA)). Anti-mouse IgG Fc Capture (AMC) biosensor tips were used to capture and immobilize antibody HB65 with increasing concentrations of rNP in the wells of solid black 96-well plates (Geiger Bio-One). The experiment was carried out in a sequential series of steps. For baselines, sensors were immersed in 1X PBS buffer for 30 seconds to obtain equilibrium and then antibody loading to sensors via movement of sensor tips to mAb HB65 antibody wells for 60 seconds in order immobilize the antibodies on the sensor tips. For the second baseline sensors were moved to wells containing 1X PBS for 300 seconds to reach equilibration. For association, sensors moved to ligand buffer (rNP samples) for 300 seconds to obtain *K*_on_. For dissociation sensors moved to 1X PBS buffer for 3,600 seconds to obtain *K*_off_. Six concentrations of ligands (rNP) were used to obtain the final curve. All the data were collected and analyzed by ForteBio (Sartorius) data analysis software. Data analysis and curve fitting of experimental data resulted in binding equations describing a 1:1 interaction. Global analysis using nonlinear least-squares fitting produced a single set of binding parameters. The equilibrium dissociation constant (*K*_D_) values were calculated using the equation (*K*_D_ = *K*_off_/*K*_on_).

### 5.8. Electron microscopy

Electron microscopy and 2D image analysis of influenza nucleoprotein (NP) and antibody samples were similar to previously reported (Gallagher et al., 2017). In brief, protein samples were incubated on a glow-discharged continuous-carbon grids, rinsed twice with water, then negatively stained with 0.25 % uranyl acetate for 1 minute, blotted with filter paper, and then allowed to dry. Grids were imaged using a Tecnai-12 cryo-electron microscope (ThermoFisher, Scientific) operating at 100 kV and images were acquired using a Gatan OneView camera (Warrendale, PA). Particles were picked and 2D classification carried with the image processing software EMAN2 (Tang et al., 2007).

## ACKNOWLEDGEMENTS

This work was supported by the Intramural Research Program of the National Institute of Allergy and Infectious Diseases, National Institutes of Health, 1ZIAAI001180, AKH. We thank Larry Lantz (NIAID) for aid in antibody production. This work utilized the computational resources of the NIH HPC Biowulf cluster (http://hpc.nih.gov). This work used reagents obtained through the Influenza Reagent Resource, Influenza Division, WHO Collaborating Center for Surveillance, Epidemiology and Control of Influenza, Centers for Disease Control and Prevention, Atlanta, GA, USA. We also would like to thank Dr. Derron A. Alves, Kevin Bock (NIAID/NIH) and Udana Torian (NCI) for input.

