## Supplemental figures for "Analysis of polyclonal and monoclonal antibody to the influenza virus nucleoprotein in different oligomeric states"

**[2] Current Address: Department of Immunology, School of Medicine, University of Washington, Seattle, WA, USA 98195**

**This PDF file includes:**

Supplemental Fig. S1 to S5

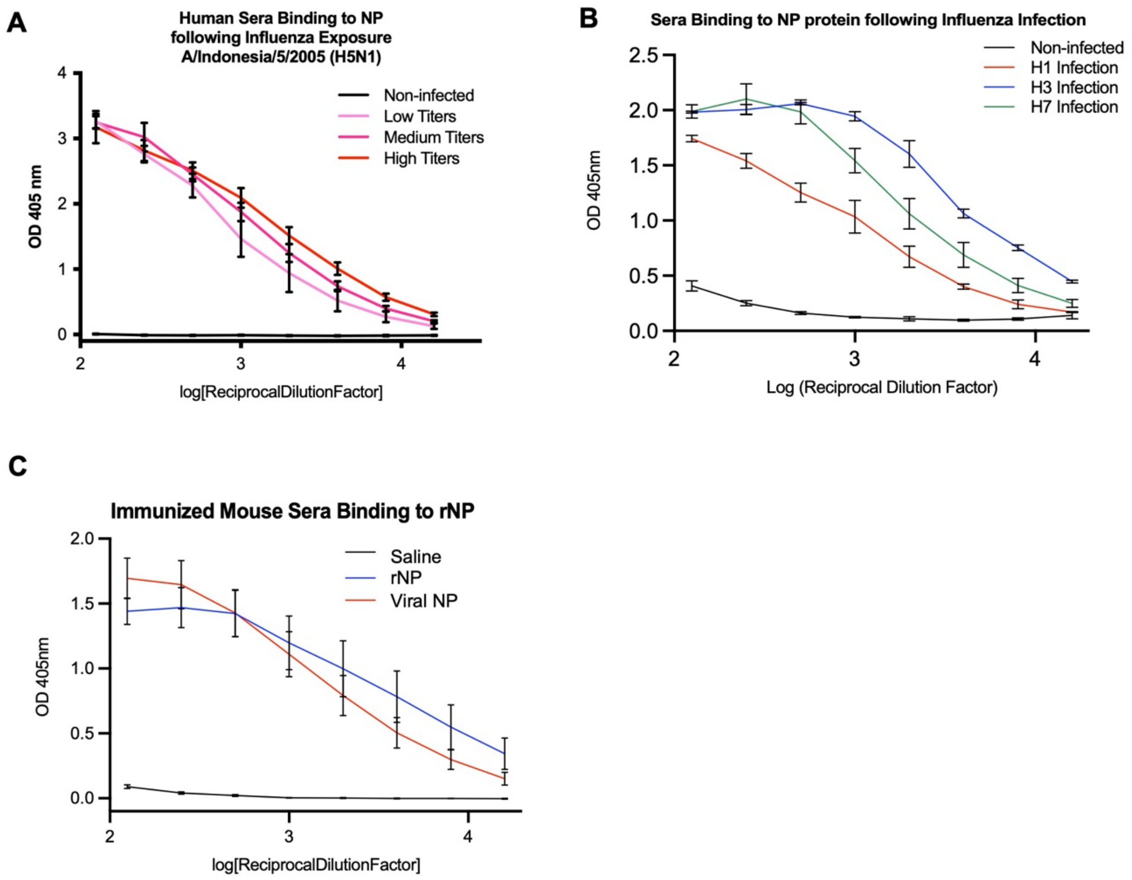


Supplemental Fig. S1

**Fig. S1. ELISA of dilution series of sera to test binding of human, ferret, and mouse sera to NP to calculate area under the curves.** (A). ELISA analysis of pooled sera from humans infected with H5N1 deemed to have low (pink), medium (magenta), and high (red) titers of virus for detection of antibody binding to rNP. (B) ELISA analysis of sera of groups of ferrets infected with H1 (red), H3 (blue), and H7 (green) influenza viruses, respectively, for detection of antibody binding the rNP. Non-infected sera were negative controls (black). (C) Probing the binding of anti-rNP (blue) and anti-viral NP (red) mouse sera to rNP. Saline was a negative control (black). Recombinant NP from influenza A/Brisbane/10/2007 (H3N2) was the antigen used for detection and ferret, human and mouse sera were used as the primary antibody, respectively. Curves were used to derive area under the curve (AUC) measurements shown in main-text Figs. 1A,1E, 2F.

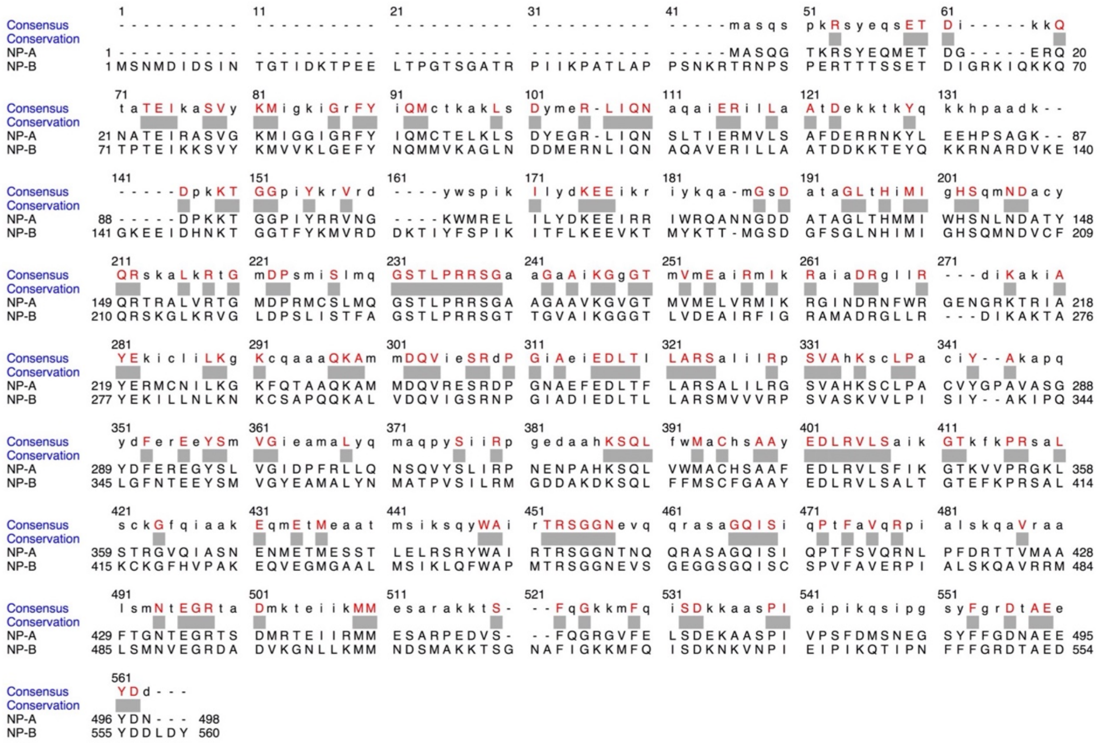

Supplemental Fig. S2

**Fig. S2. Sequence alignment of nucleoproteins (NP) from a type A (NP-A) and a type B (NP-B) influenza virus.** NP-A sequence is of influenza A/Puerto Rico/8/1934 (H1N1) and NP-B sequence is of B/Lee/40. A consensus conservation sequence is shown above the two aligned sequences with identical residues colored red and indicated by grey boxes.

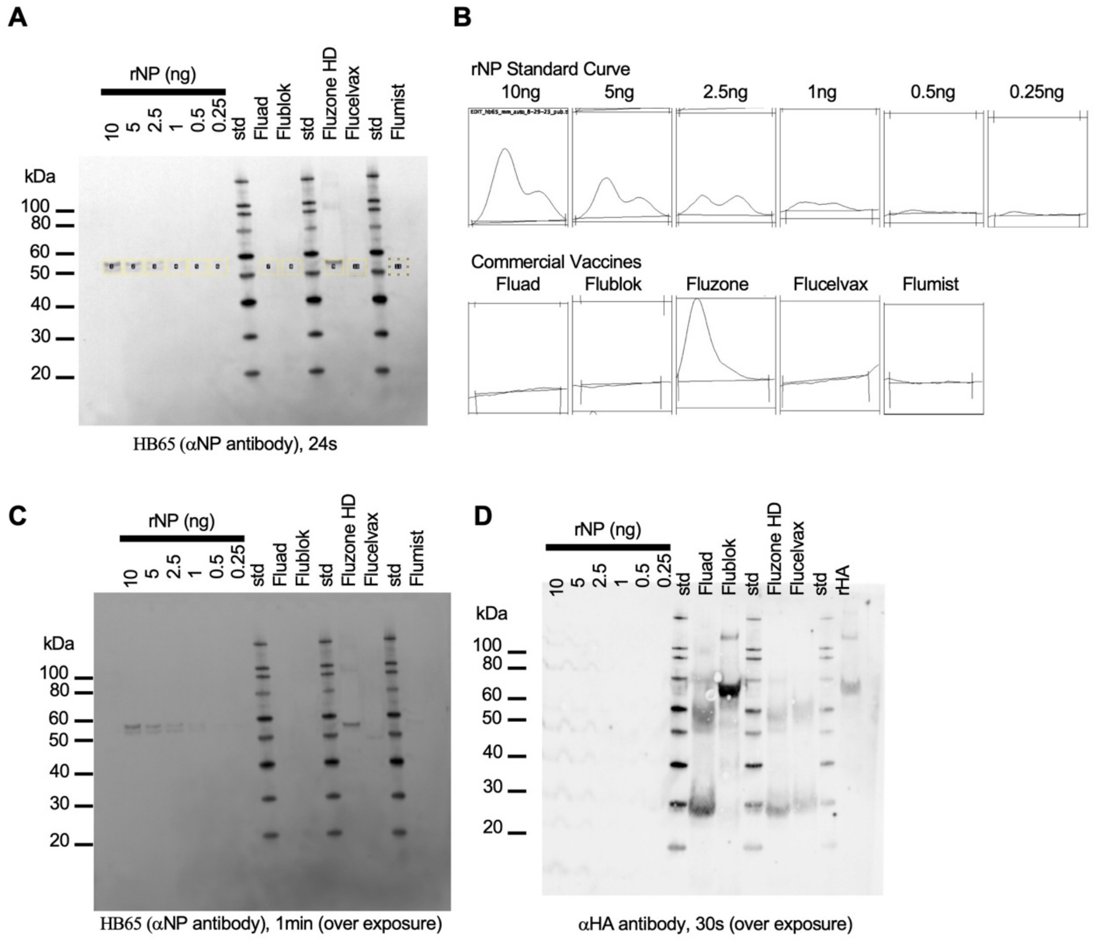


Supplemental Fig. S3

**Fig. S3. Quantitation of NP in commercial influenza vaccines using anti-NP mAb HB65 and recombinant NP.** (A) Immunoblot with samples consisting of deceasing amounts of rNP and different commercial influenza vaccines. Regions used in quantitation are boxed (yellow). (B) Corresponding density profile traces from boxed areas in panel A. (C) Immunoblot in panel A with increased exposure time. (D) Protein transfer confirmation to blot via probing with anti HA antibody to detect HA in commercial influenza vaccines. Commercial influenza vaccines were Flublok, Fluvirin, Fluzone HD, and Flucelvax. Standards are denoted (kDa). Recombinant NP was from influenza A/Brisbane/10/2007 (H3N2).

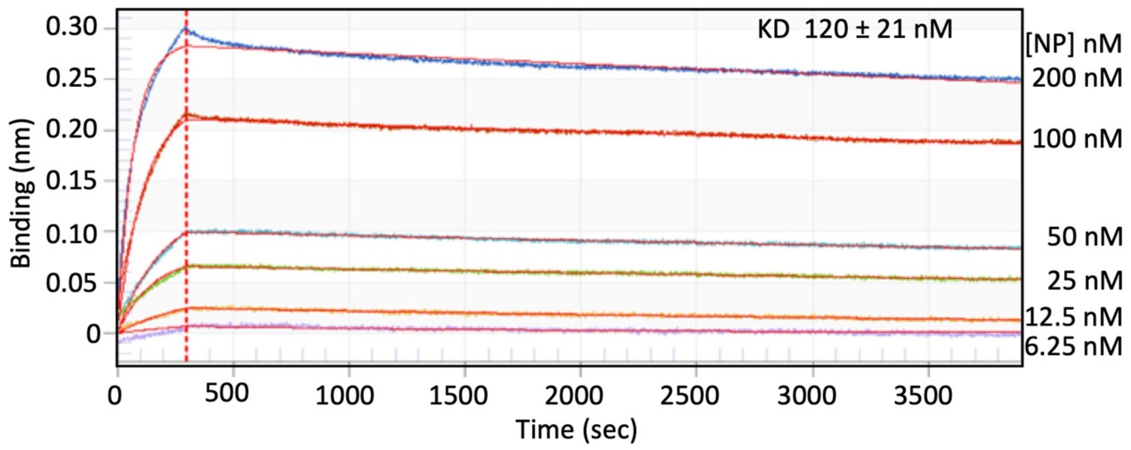


### Supplemental Fig. S4

**Fig. S4. Measurement of the affinity between anti-NP mAb HB65 and nucleoprotein (NP) using biolayer interferometry.** Octet sensograms for measurement of anti-NP mAb HB65 binding to NP by bio-layer interferometry (BLI). NP was used as analyte in solution and mAb HB65 was immobilized onto the bait probe (Anti-Mouse Fc Capture Biosensor). NP was the analyte in solutions with varying concentrations (6.25-200 nM). Curve fits are in red. Binding is measured in BLI by an increase in optical thickness of the bait probe that results in a wavelength shift that is measured in nanometers. The measured $K_D$ was 120 ± 21 nM.

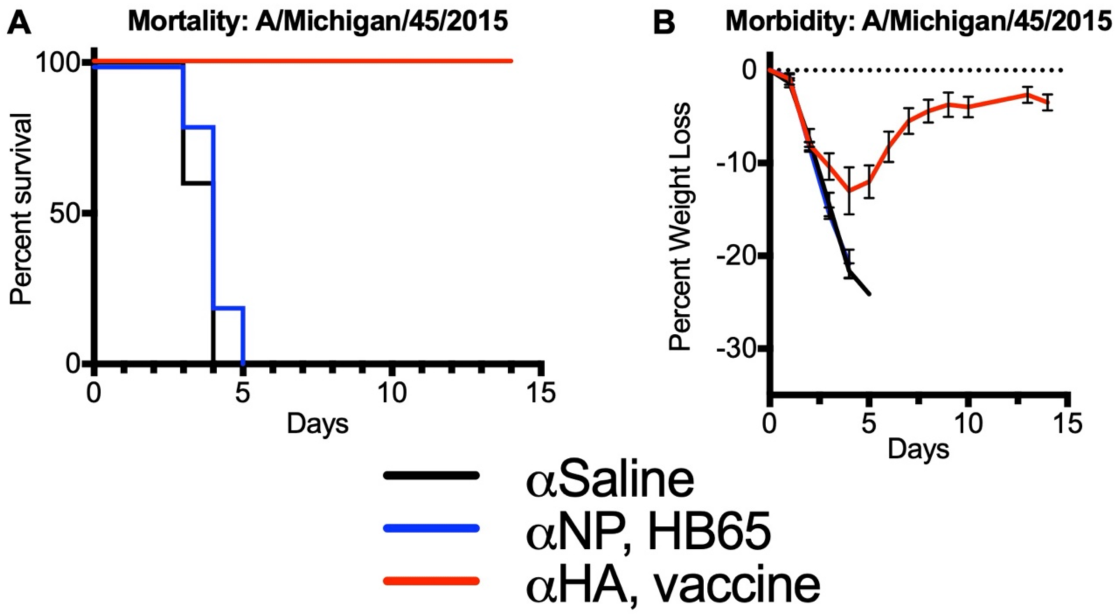


Supplemental Fig. S5

**Fig. S5. Passive-transfer of anti-NP mAb HB65 and vaccine sera for protection from H1N1 challenge**. (A) Survival curves for mice that were intranasally challenged with influenza virus (A/Michigan/45/2015 (H1N1)) 18 hours after intraperitoneally (IP) transfer of 400ul of mAb HB65 (0.875mg/ml) (blue line) or sera collected from mice immunized with commercial influenza vaccine (Flublok) (red line). (B) The corresponding weight-lost curves for panel A, with mAb HB65 (blue line), positive control Flublok vaccine sera (red line) and saline negative-control (black line).
